# Pharmacological mechanisms on the anti-breast cancer property of *Coix lacryma-jobi*: A network-based analysis

**DOI:** 10.1101/2021.10.16.464646

**Authors:** Angelu Mae Ferrer, Janella Rayne David, Arvin Taquiqui, Arci Bautista, Custer C. Deocaris, Malona V. Alinsug

**Affiliations:** Department of Biology, College of Science, Polytechnic University of the Philippines, Manila, Philippines; Atomic Research Division, Department of Science & Technology – Philippine Nuclear Research Institute, Central Avenue, Diliman, Quezon City, Philippines

**Keywords:** Bioactive compounds, *Coix lacryma-jobi*, network pharmacology, targets, breast cancer

## Abstract

Breast cancer is considered as one of the three most common cancers around the world and the second leading cause of cancer deaths among women. *Coix lachrymal jobi,* commonly known as Job’s tears or adlay has been reported to possess anti-cancer properties. Despite evidences provided by clinical data, the usage of *Coix lacryma-jobi* in treating cancer, particularly breast cancer, has been scarce. Thus, this study was conducted to determine the pharmacological mechanisms underlying its anti-breast cancer property using various network pathway analyses. Bioactive compounds from *Coix lacryma-jobi* and its potential target genes were obtained from SymMap. Breast cancer-related target genes were collected from CTD. Protein-protein interaction network was analyzed using the STRING database. GO and KEGG pathway enrichment analyses were performed using DAVID to further explore the mechanisms of *Coix lacryma-jobi* in treating breast cancer. PPI and compound-target-pathway were visualized using Cytoscape. A total of 26 bioactive compounds, 201 corresponding targets, 36625 breast cancer-associated targets were obtained, and 200 common targets were considered potential therapeutic targets. The 9 bioactive compounds identified were berberine, oleic acid, beta-sitosterol, sitosterol, linoleic acid, berberrubine, jatrorrhizine, thalifendine, and stigmasterol. The identified 5 core targets were ESR1, JUN, MAPK14, and RXRA. *Coix lacryma-jobi* targets enriched in GO terms were mostly involved in regulation of transcription from RNA polymerase II promoter, drug response, steroid hormone receptor activity, and protein binding. This study elucidates on the pharmacological underpinnings on the potency of adlay against breast cancer. Its subsequent drug development will be worth a step forward for a breast cancer-free society.

## INTRODUCTION

Breast cancer arises from the epithelial lining of the ducts or globules in the glandular breast tissue. According to World Health Organization, breast cancer is one of the most frequent cancer affecting women with approximately 685,000 deaths reported in 2020. Every year, about 1.5 million women worldwide (25% of all cancer patients) are diagnosed with breast cancer. It is a metastatic cancer that can spread to distant organs including the bone, liver, lung, and brain which explains the difficulty in treating this disease. However, an excellent prognosis and a high survival percentage can be achieved if the disease is detected early (Sun et al., 2017).

Although modern medication is primarily prescribed for treatment of breast cancer, traditional herbal medicine is used as supplemental and alternative method in several developed countries (Abu Bakar et al., 2021). The use of chemotherapeutic agents for its treatment are of plant origin like leaves, fruits, flowers, fungi, and lichens. The term “herb” refers to plants that produce fruits and seeds. These herbs are important as they are used to maintain good health (Shareef et al., 2016).

*Coix lacryma-jobi*, also known as Job’s tears or adlay is widely cultivated in China and Japan as a nutritious food supplement. Proteins, essential amino acids, and carbohydrates are just some of the nutrients found in Coix seeds. In addition, many important classes of bioactive compounds found in Coix seeds including coixenolide, triglyceride, fatty acids, and triterpenes, have been found to provide numerous health benefits (Yu et al., 2017).

Adlay has been found to have an anti-tumor activity in several studies. Berberine, one of its bioactive compounds, is observed to suppress tumors, thus preventing cancer (Manosroi et al., 2016). Past studies have also shown that Coix seed extract has anti-inflammatory, antioxidative stress, and anti-cancer properties (Hien et al., 2016). Furthermore, the Chinese Ministry of Public Health has authorized the emulsion of Job’s tears oil for anti-cancer action. Patients have used this medicine to treat different types of cancers, such as lung, breast, and liver cancer (Kuo et al., 2016).

Having these observations in *Coix lacryma-jobi* containing anti-cancer properties, this study assessed its potential role in the treatment of breast cancer using a network pharmacological approach. The study evaluated the interactions between the bioactive compound targets of *Coix lacyrma-jobi* L. and disease targets of breast cancer. The mechanism of the bioactive compounds that may have therapeutic mechanism for breast cancer was also explored.

## MATERIALS & METHODS

### 2.1 Information acquisition of compound composition of Coix lacryma-jobi and breast cancer

The compound composition of coix seed were retrieved from the ‘ingredient tab’ in SymMap (http://www.symmap.org/, Version 2), a database that provides massive information on herbs, ingredients, targets, as well as clinical symptoms and diseases (Wu et al., 2019). Data acquisition of *Coix lacryma-jobi* was done using *‘Coix lacryma-jobi’* as the keyword. Other information of the obtained components including, molecular weight, PubChem ID, and canonical smiles were confirmed and obtained using PubChem database (https://pubchem.ncbi.nlm.nih.gov/). To construct a network with breast cancer, target genes of the disease with the *“Cancer of Breast”* as the keyword were searched and obtained from the Comparative Toxicogenomics Database (CTD, https://ctdbase.org/, updated 2^nd^ September 2021) through the ‘gene’ tab and with limitations to disease identifier. CTD is a manually curated information about the integrated interactions of chemicals, genes, phenotypes, diseases, and how environmental exposures affects human health (Davis et al., 2020). Information obtained from these databases were imported and organized in separate sheets to avoid mix-ups of data and were saved as TSV files.

### 2.2 Acquisition of Bioactive Compounds through Pharmacokinetic ADME Evaluation

All retrieved compounds were screened with oral bioavailability (OB) value and druglikeness (DL). Initially, compounds were retained only if OB ≥ 30 (Wang et al., 2017). Compounds without OB information were also removed from the list. Then, using the Swiss ADME website (http://www.swissadme.ch/) the retained compounds were screened with DL with limitation to the Lipinski’s Rule of Five (RO5) by inputting the Canonical SMILES corresponding to each compound. A compound with more than one violation were removed from the list. Compounds that satisfied the criteria were identified as potential bioactive compounds.

### 2.3 Intersection of bioactive compound target genes and disease related target genes

The obtained *Coix lacryma-jobi-related* target genes and breast cancer-associated target genes were separately imported into Microsoft (MS) excel spreadsheet to remove duplicate genes. Both target genes were combined into a single column and duplicates were screened and copied into a different sheet. From that, duplicates were then removed to obtain the common target genes.

### 2.4 Protein-Protein Interaction (PPI) Network Analysis

The common target genes of *Coix lacyrma-jobi* and breast cancer were uploaded to Search Tool for the Retrieval of Interacting Genes (STRING, https://string-db.org/, Version 11.5) as the background network database with limitation to *“Homo sapiens”* and a highest confidence score of 0.900 to construct a high-quality and credible PPI network (Zhou et al., 2021). The target protein interaction was obtained and saved as a TSV file. Then the TSV file was imported into Cytoscape software (http://www.cytoscape.org/, ver 3.8.2) to analyze and visualize the PPI network. Using the Network Analyzer tool, summary statistics of the nodes and edges, along with the topological parameters such betweenness centrality, closeness centrality, and the degree values of each target were obtained.

### 2.5 Enrichment Analysis and Network Construction

The Database for Annotation, Visualization, and Integrated Discovery (DAVID, https://david.ncifcrf.gov/home.jsp, version 6.8) was used for GO and KEGG pathway functional enrichment analysis. By starting the analysis, the Gene symbol of common targets of “CS-BC” was imported in the gene list. OFFICIAL_GENE_SYMBOL, *Homo sapiens,* and gene list were selected in the identifier, species, and list type, respectively. The screening criteria was limited to P ≤0.05 using Bonferonni correction (Liang et al., 2014). Biological process (BP), Cellular Component (CC), and Molecular Function (MF) are the three components of GO term enrichment analysis. GOTERM_BP_DIRECT, GOTERM_CC_DIRECT, and GOTERM_MF_DIRECT, as well as KEGG_PATHWAY were selected in Gene Ontology and Pathways in the annotation category, respectively to analyze and visualize the results. Pathways with the top 20 protein numbers were used for the establishment of the compound-target-pathway network by Cytoscape. Figure 1 presents the overall workflow of the methods conducted in this study.

**Figure 1.**
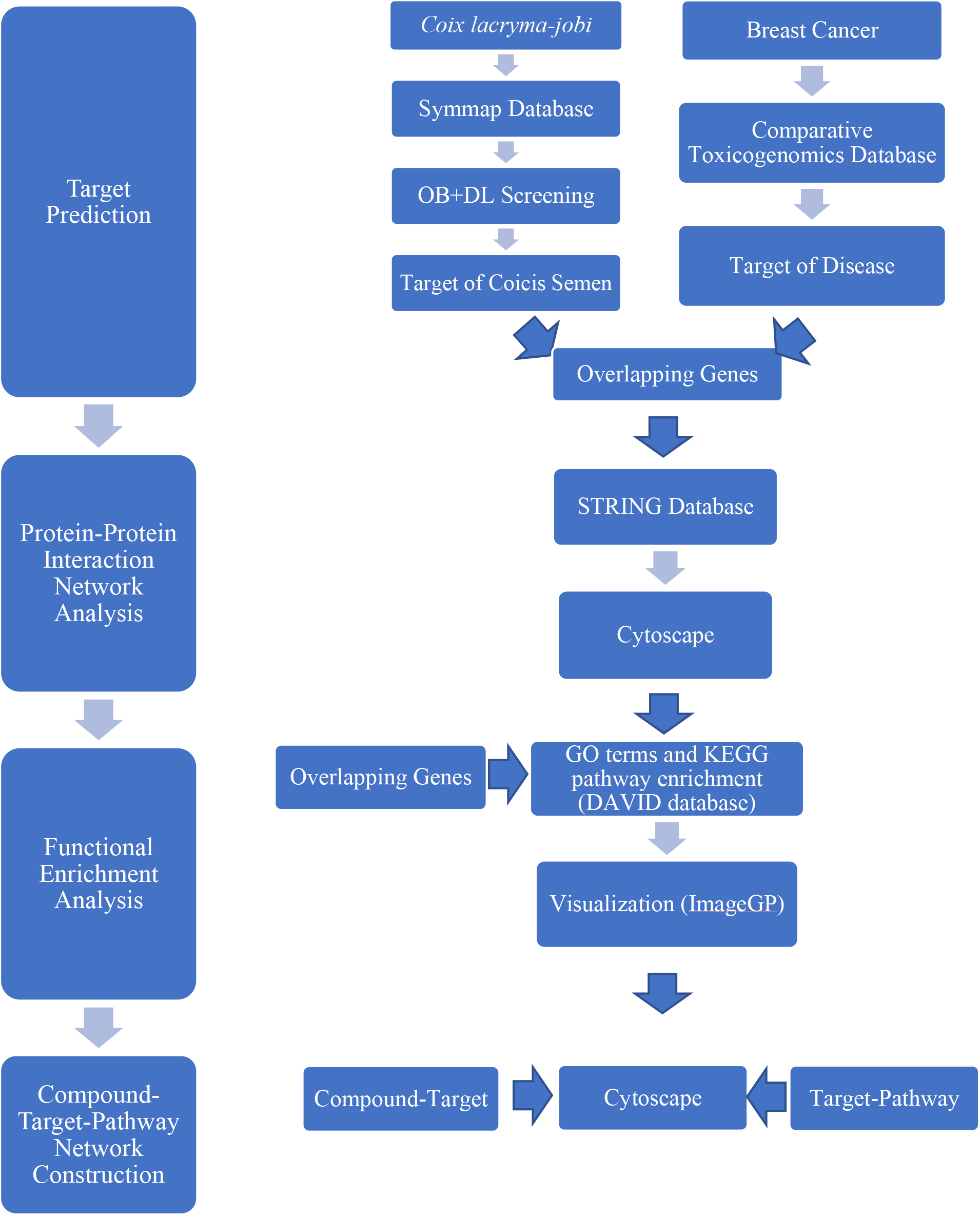
Schematic Workflow. Summary of methods conducted for the network analysis of *Coix lacryma-jobi* components against breast cancer is presented.

## RESULTS

### 3.1 Compounds, Corresponding Targets, and Genes of *Coix lacryma-jobi*

There were 111 compounds of *Coix lacryma-jobi* obtained from SymMap database. So far, 113,399 target genes were related to breast cancer from CTD. A total of 507 were identified as potential target genes of adlay from SymMap.

### 3.2 Collection of Bioactive Compounds and Targets of *Coix lacryma-jobi* using ADME Evaluation

Drug treatments are usually taken orally for clinical medication (Li et al., 2021). Thus, it is essential that from the selection up to identification to be an effective treatment for the patient, a compound must exhibit high biological activity and low toxicity (Daina et al., 2017). Daina et al. (2017) claimed that the initial assessment of ADME (Absorption, Distribution, Metabolism, and Excretion) in the discovery phase exhibited a significant reduction of the failures associated with pharmacokinetics in the clinical phases. In keeping with the traditional ADME screening principle (Xu et al., 2020), oral bioavailability and drug likeness are the two main variables used for the screening of compounds that affect drug absorption across the gastrointestinal mucosa (Huang et al., 2020).

Oral bioavailability (OB) is the intensity and rate of absorbing drugs into the systemic circulation. The value of the percentage of the OB of the compound is directly proportional to the probability of the clinical application development of the compound (Chen et al., 2018). High OB is normally a core indicator in the establishment of the drug likeness index of bioactive molecules (Huang et al., 2020). Drug-likeness (DL) represents the capacity of molecules in the compound demonstrating physicochemical properties comparable to existing drugs (Potunuru et al., 2019). Lipinski’s Rule of Five (RO5) is an empirical guideline that predicts the high risk of poor absorption or permeability of a compound on its underlying simple physicochemical properties. For a compound to be considered an orally available drug, it must satisfy the following conditions: (1) no more than 5 hydrogen-bond donors, (2) no more than 10 hydrogen-bond acceptors, (3) the molecular weight (MWT) is less than 500 Dalton, and (4) the calculated Log P (CLogP) or lipophilicity is less than 5 (A decade of Drug likeness, 2017). A compound with 2 or more violations of these conditions is considered to be non-orally available drug (Lipinski et al., 2001).

The screening condition for OB was set as ? 30 (Chen et al., 2018). For DL, a compound must have no more than 1 violation from the Swiss ADME. A compound that satisfied both criteria was considered a bioactive compound. From the 111 compounds retrieved from SymMap, 30 were with oral bioavailability greater than or equal to 30. 2 of the compounds were unable to be found in PubChem and have no PubChem ID while 1 have no target genes. Finally, after ADME evaluation and potential target gene prediction, a total of 25 compounds were identified as potential bioactive. Detailed information on *Coix lacryma-jobi* compounds is presented in Supplementary Tables S1 and S2.

The corresponding target genes of each identified bioactive compound were obtained from SymMap. From the 507 target genes, a total of 201 target genes were obtained after the removal of duplicates. Detailed information on *Coix lacryma-jobi-related* targets is presented in supplementary Table S3.

### 3.3 Identification of Targets Related to Breast Cancer

CTD was mainly used to identify target genes related to breast cancer. From 113,399 potential related disease targets genes, a total of 36,625 target genes associated with breast cancer were identified after the removal of duplicate targets. Detailed information on DM-related targets is presented in Supplementary Table S4.

The bioactive compound targets and disease targets were intersected to obtain 201 common targets as shown in Figure 2. These common targets were taken as the central target for subsequent enrichment analysis and protein-protein interaction network.

**Figure 2.**
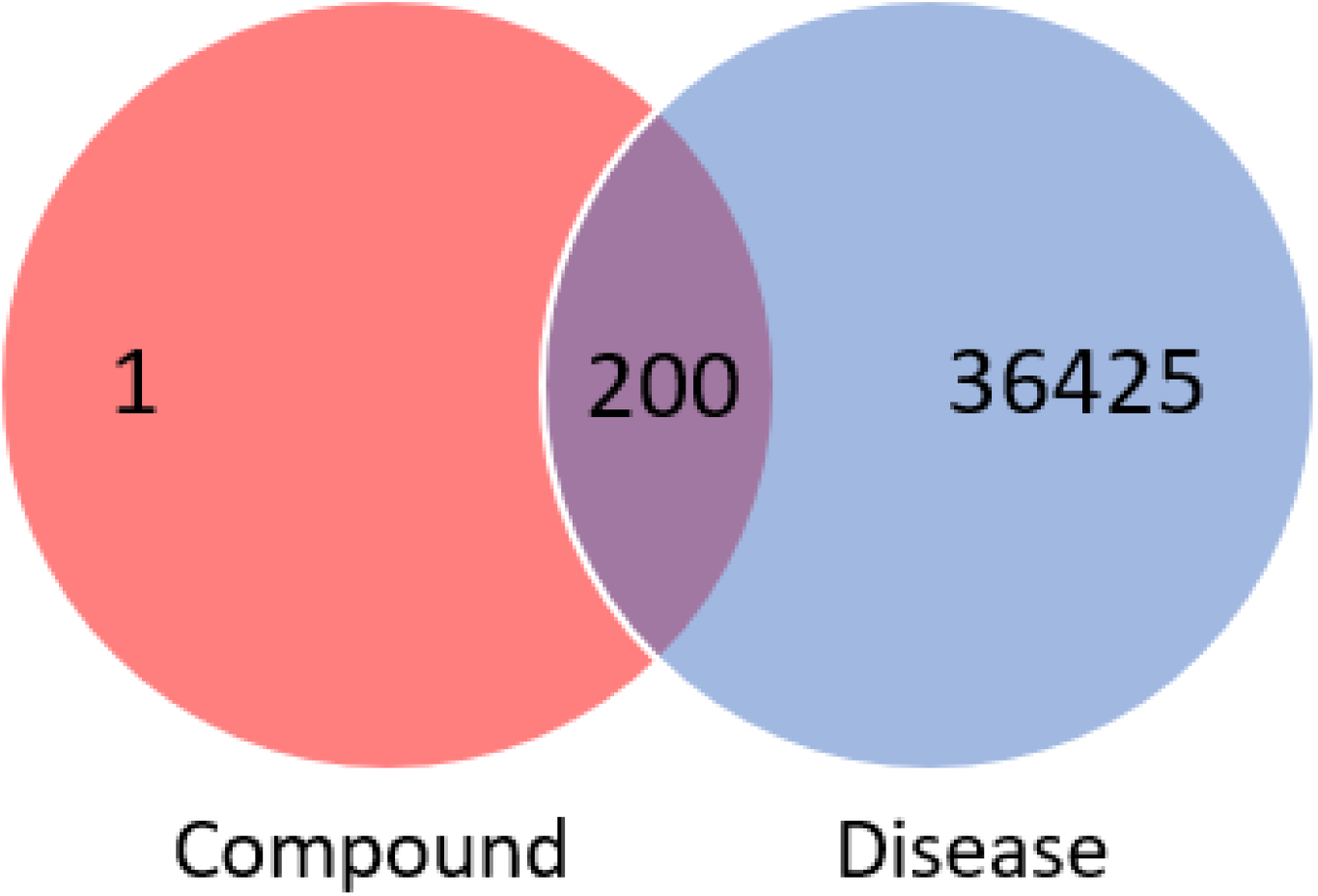
Venn diagram of *Coix lacryma-jobi* compound targets and breast cancer disease

### 3.4. Protein-Protein Interaction Network

The PPI analysis of 201 common targets of *Coix lacryma-jobi* and breast cancer was executed through String database with limitation to *“Homo Sapiens”* and with a confidence score of 0.900. The PPI network then was established and visualized using Cytoscape 3.8.2, of which results (Figures 3–4) generated a total of 153 nodes representing all the predicted targets; with 652 edges representing the correlation between the active components of *Coix lacryma-jobi* and potential breast cancer targets. Network Analyzer tool in Cytoscape was utilized for the calculation of the topological parameters of the PPI network to identify hub nodes and essential targets wherein nodes with higher betweenness centrality were depicted by darker color in the network. Based on the result from the network analyzer, the top 10 targets of coicis-breast cancer are MAPK3, JUN, MAPK1, TP53, RXRA, RELA, NCOA1, MAPK14, ESR1 and FOS as defined by the highest degree and were described by a larger size in the network. These were then identified as hub nodes and essential targets of the PPI network.

**Figure 3.**
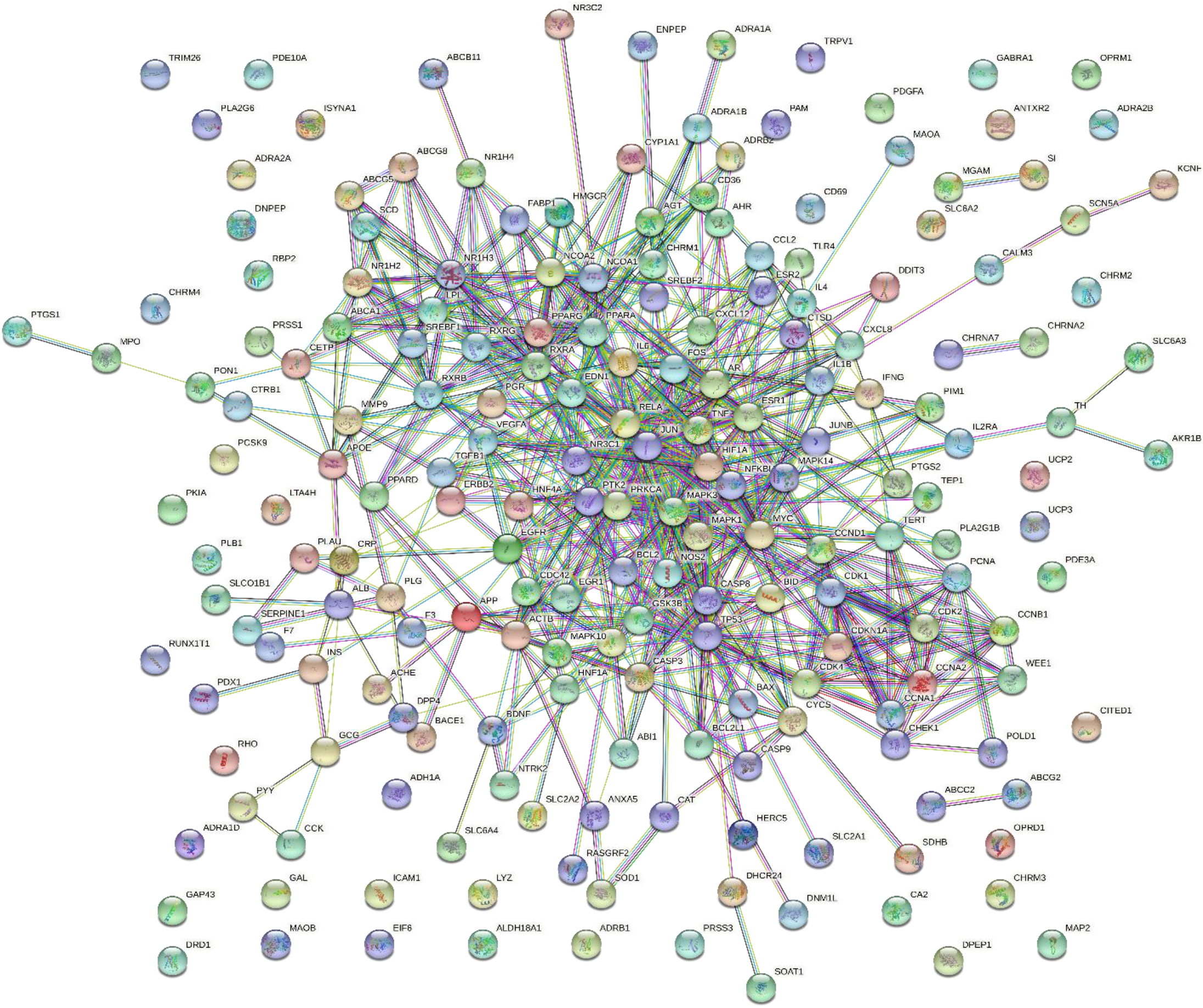
Interaction relationship between target proteins of *Coix lacryma-jobi* in breast cancer was analyzed using STRING. PPI enrichment was set at a p value <1.0e-16.

**Figure 4.**
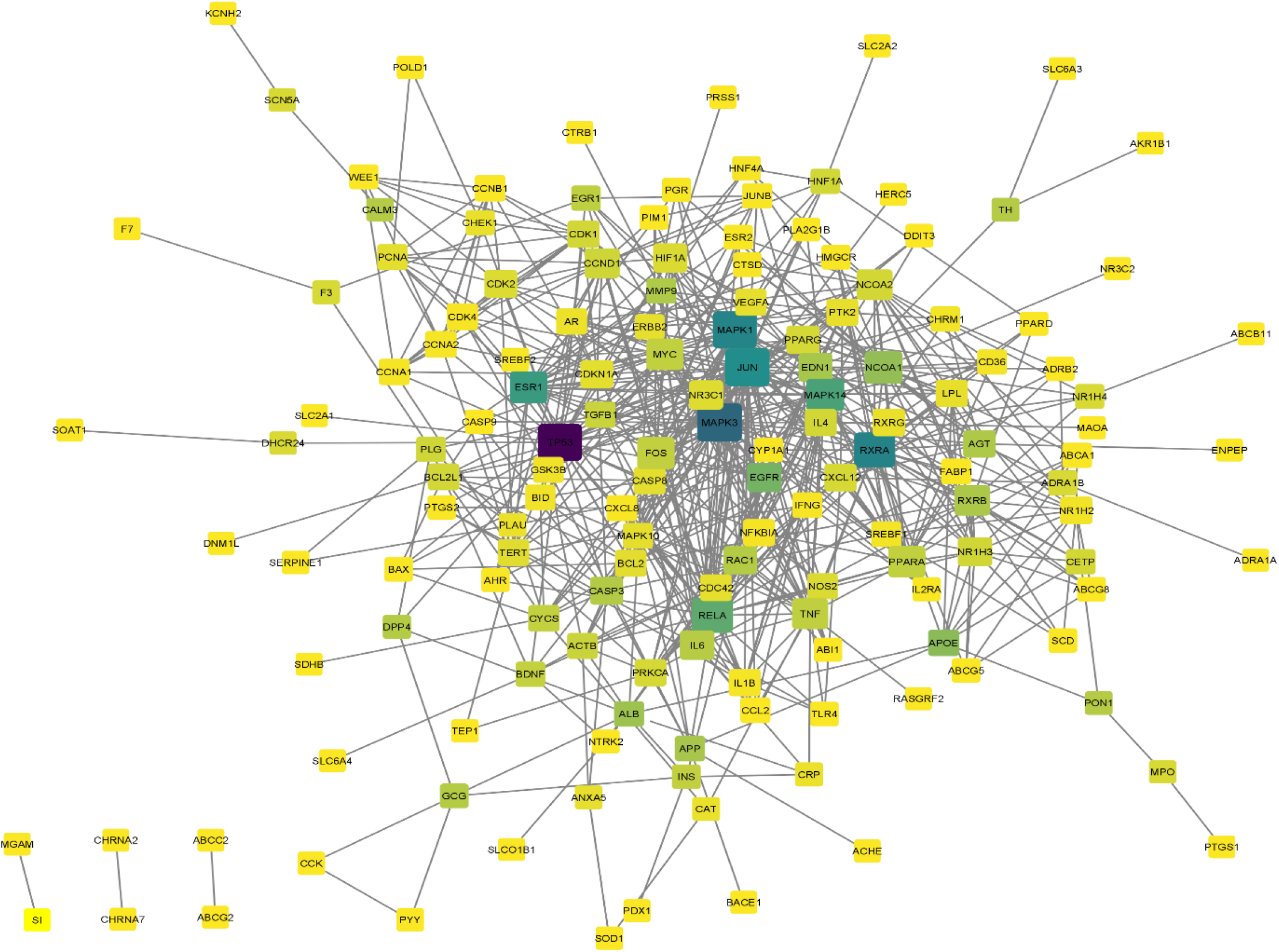
Interaction network of potential proteins of *Coix lacryma-jobi* in treating breast cancer was based on Cytoscape composition results.

### 3.5. GO and KEGG Pathway Enrichment Analysis

The GO and KEGG Pathway Enrichment Analysis using the 201 common targets of *Coix lacryma-jobi* and breast cancer was executed through the DAVID database to identify potential mechanisms of *Coix lacryma-jobi* against breast cancer with a p-value cut-off of 0.05 as the screening condition. Bonferroni values were used for the identification of the top 20 significantly enriched GO terms and KEGG pathways of the gene overlaps (Figure 5). Biological Process (BP), Cell Component (CC) and Molecular Function (MF) were used as the main components of the GO term enrichment analysis and KEGG for the pathway Enrichment Analysis. According to the results, the 200 targets were significantly enriched in 500 BPs, 64 CCs, 109 MFs and 114 KEGG pathways. In the biological process (BP) category, the target proteins were mainly involved in response to drug, positive regulation of transcription from RNA polymerase II promoter and negative regulation of apoptotic process. In the cellular component (CC) category, the target proteins were mainly involved in cytoplasm, plasma membrane and extracellular space. Molecular Function (MF) analysis has a higher enrichment of steroid hormone receptor activity, enzyme binding and protein binding.

**Figure 5.**
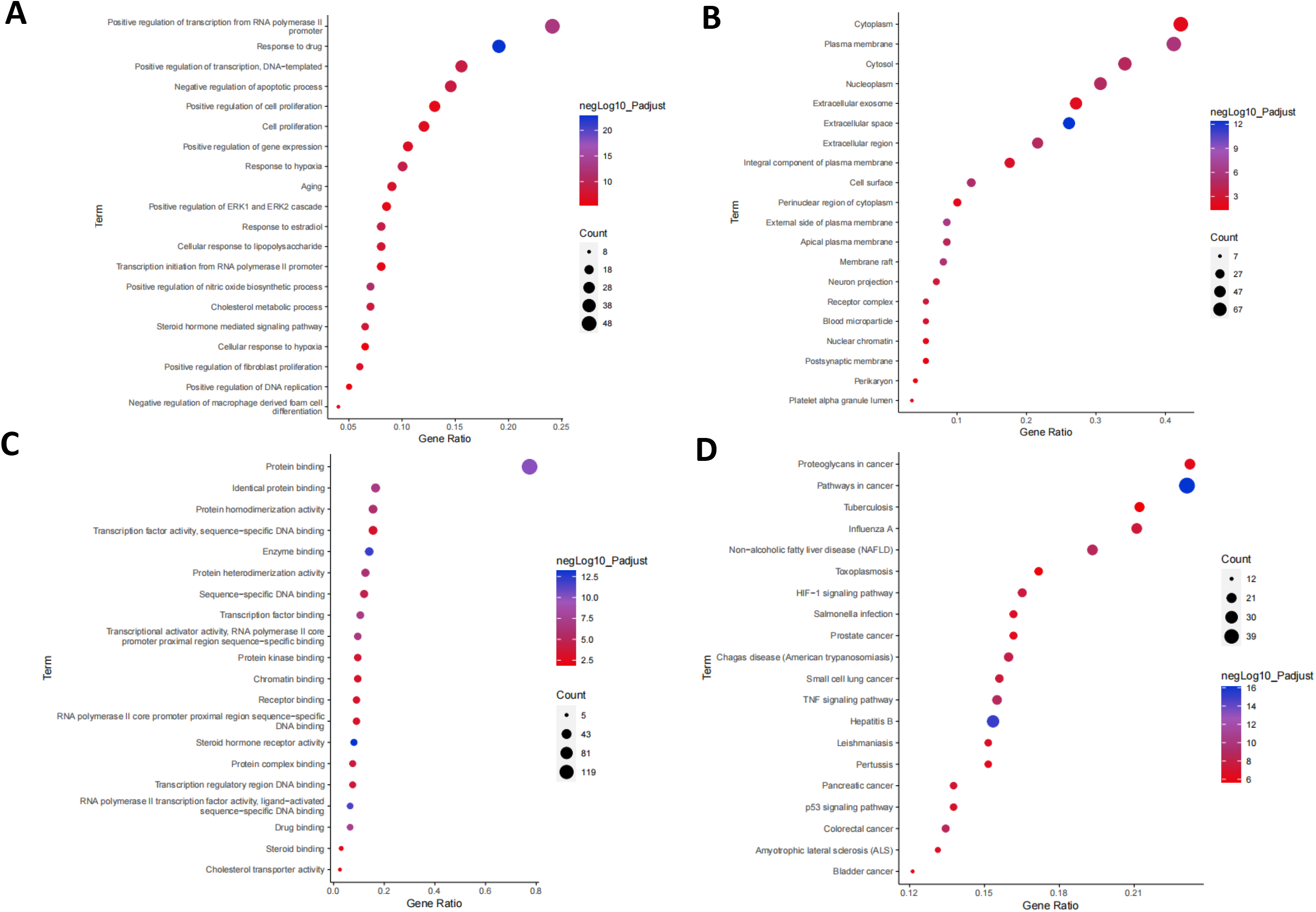
Pathway Analysis. GO analyses and KEGG pathways enrichment results were based on the following: (A) Biological Processes; (B) Cellular Components; (C) Molecular Functions; (D) KEGG pathways.

**Figure 6.**
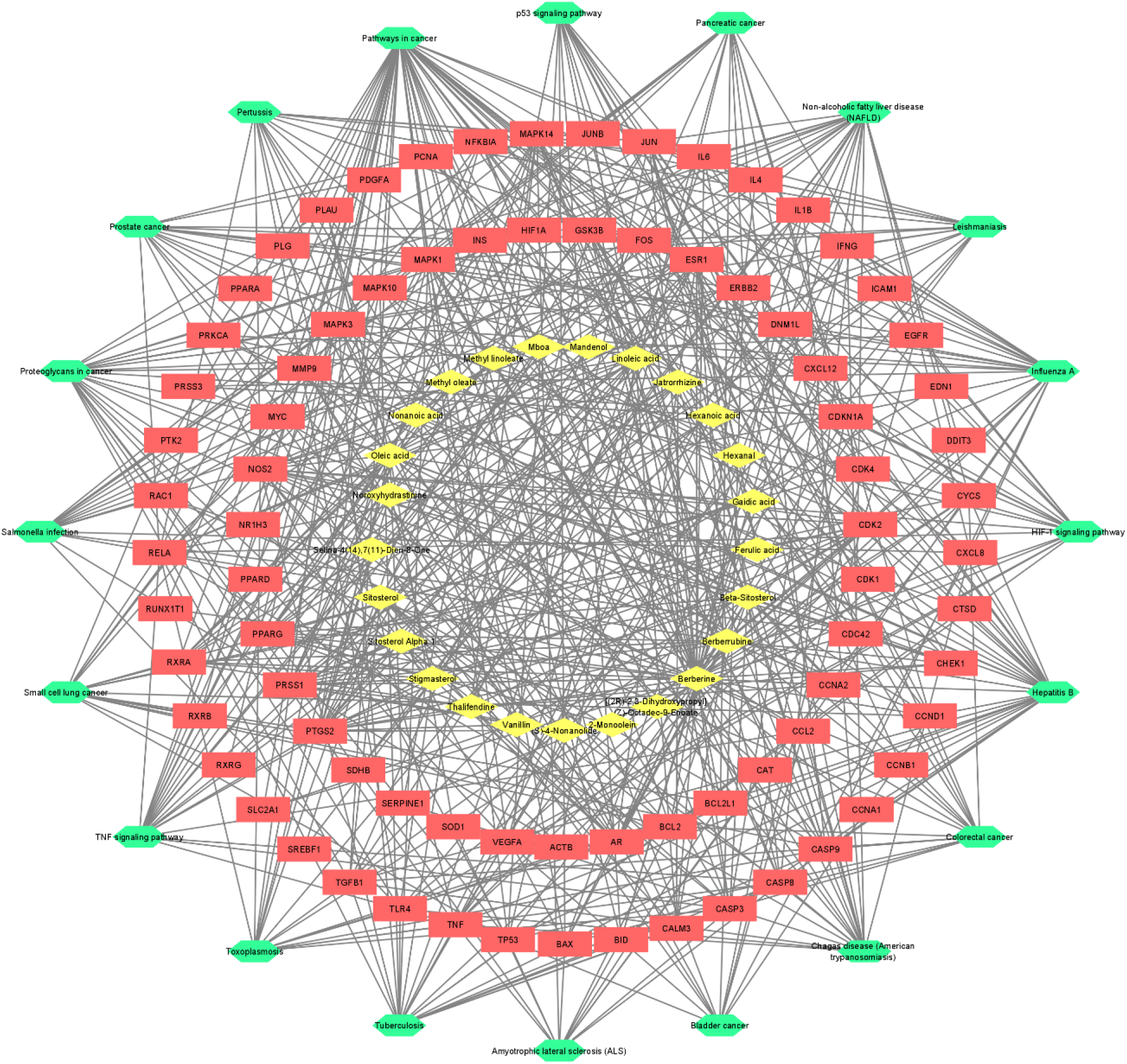
Compound-Target-Target Pathway Network. Yellow diamonds represent bioactive compounds of *Coix lacryma-jobi,* red rectangles represent target genes, and green hexagons represent the top 20 pathways.

The KEGG enrichment analysis results showed that pathways in cancer was the most closely related to the function of adlay in breast cancer.

### 3.6. Construction of Compound-Target-Target Pathway Network

The constructed compound-target-pathway network was established using Cytoscape 3.8.2 according to KEGG pathway enrichment result. The compound-target-pathway included 125 nodes and 607 edges, yellow diamonds represent bioactive compounds of *Coix lacryma-jobi*, red rectangles represent target genes, and green hexagons represent the top 20 pathway. Edges represent the relationship between compound and target, and between target and pathways. Hence, targets are bridges between compounds and pathways. In the network, 9 compounds had a higher-than-average degree, which revealed that they performed a key role in the compound-target-pathway network. The compounds were Berberine, Oleic acid, Beta-Sitosterol, Sitosterol, Linoleic acid, Berberrubine, Jatrorrhizine, Thalifendine, Stigmasterol. The top 8 hub nodes and essential targets of the PPI network were identified based on the overlap of the top 20 targets of degree, between centrality, and closeness centrality, which were ESR1, JUN, MAPK1, MAPK14, MAPK3, RELA, RXRA, and TP53. The interaction of the top 8 targets of the PPI network and the compound-target-pathway network was identified as core targets, indicating that they play a significant role in both PPI network and compoundtarget-pathway network. Lastly, four targets, ESR1, JUN, MAPK14, and RXRA were identified as core targets.

## DISCUSSION

Breast cancer is considered to be one of the three most common cancers around the world (Harbeck & Gnant, 2017). In fact, it is the second leading cause of cancer deaths among women (Sun et al., 2017). It is described as a disease that begins in the breast cells that go out of control. BC commonly starts and accumulates in different parts of the breast including the ducts and lobules that will eventually metastasized to other parts of the body (CDC, 2021.). In male, breast cancer is a rare condition (Darre et al., 2020) that accounts for less than 1% of all cancers in men breast (Giordano et al., 2018) and less than 1% of all diagnosed cancers (Perkins, 2003). However, an escalation of number of breast cancer-diagnosed men have been reported, although at a lower rate (De Carmen et al., 2021). Generally, breast cancer poses high morbidity and mortality rate worldwide and cause serious health threat (Fang et al., 2020). At present, treatments for BC mainly includes surgery, radiotherapy, hormone therapy, targeted therapy, and chemotherapy (Akram et al., 2017). However, frequent and long-term use of these treatment methods especially chemotherapy often induce adverse side effects. As to why use of herbal medicines are being looked upon, with various herbs being studied due to their promising therapeutic potentials in improving the clinical treatment of diseases such as breast cancer with reduced toxicity and minimal side effects. In this study, the active constituents of *Coix lacryma-jobi* and its therapeutic potential for the treatment of breast cancer were explored in a network-based approach. From the SymMap database, a total of 201 bioactive chemicals of adlay were collected for this study based on multiple screens. Following that, breast cancer-associated genes collected from the CTD database were intersected with the targets of the bioactive compounds of *Coix lacryma-jobi,* of which generated a total of 200 final target genes that were used for the interaction and pathway analysis in this study.

Based on the results of the PPI network analysis, constituents of *Coix lacryma-jobi* are closely related to various target proteins in breast cancer inferring that the therapeutic efficacy of *Coix lacryma-jobi* in the treatment of breast cancer relies on the combined action of several target proteins rather the regulation of a single target protein. With this, the top 10 targets of coix-breast cancer with the highest degree based on the PPI network were selected which include TP53, JUN, RELA, MAPK14, NCOA1, FOS1, MAPK3, MAPK1, RXRA, and ESR1.

Moreover, the enrichment results revealed that 500 BPs, 64 CCs, 109 MFs and 114 pathways were involved. Nine (9) bioactive compounds and five (5) core targets were identified based on the topological parameters of the PPI network and compound-target-pathway network. The 9 bioactive compounds were berberine, oleic acid, beta-sitosterol, sitosterol, linoleic acid, berberrubine, jatrorrhizine, thalifendine, and stigmasterol. The 5 core targets were identified as ESR1, JUN, MAPK14, and RXRA which could play a crucial role in the pathway of breast cancer treatment. Pathways in cancer, proteoglycans in cancer, TNF signaling pathway, colorectal disease, and small cell lung cancer were among the pathways being targeted by these genes, which are most closely related to the function of *Coix lacryma-jobi* in treating breast cancer. Meaning, *Coix lacryma-jobi* may exert properties in the treatment of breast cancer through these targets.

The nuclear receptor coactivator 1 (NCOA1) interact with nuclear hormone receptors and other transcription factors (TFs) to facilitate the assembly of transcriptional protein complexes for chromatin remodeling and activation of gene expression (Qin et al., 2015; Qin et al., 2014). It performs crucial functions in development, growth, reproduction and metabolism, as well as in cancer (Qin et al., 2015). NCOA1 has previously been reported to be overexpressed in breast cancer and its increased expression positively correlated with disease recurrence and metastasis, hence a major regulator in breast cancer (Ma et al., 2017; Qin et al., 2015; Lui et al., 2019). Consequently, studies recommend that targeting overexpressed NCOA1 in certainly a sufficient strategy to regulate growth of breast cancer and/or metastasis (Qin et al., 2015).

The mitogen-activated protein kinase 14 (MAPK14) or p38α, is a member of MAPK signaling pathway that performs an essential role in cell integration for several biochemical signals, and are involved in multiple cellular process including proliferation, differentiation, transcription regulation and development (NCBI, 2021; Liu et al., 2019). Also, MAPK14 has been described as a significant part in the coordination of DNA damage response and regulating instability of chromosome during the progression of breast cancer, including inflammatory disorders, neurodegenerative diseases, and cardiovascular cases. Thus, MAPK14 promotes the occurrence and progression of breast cancer through the activation of its downstream target genes. The decrease in level of MAPK14 results to the damage of DNA and rise in instability of breast cancer cells, eventually causing the apoptosis of cancer cells tumor deterioration. Consequently, inhibitors of MAPK14 can be used for treating breast cancer (Lui et al., 2020; Lui et al., 2019; Madkour, 2021).

The ESR1 encodes the human estrogen receptor α (ERα) which is a member of the steroid/nuclear receptor superfamily and functions as ligand-activated transcription factors (Boldes et al., 2020) involved in cell proliferation and activation (Zundelevich et al., 2020) and have been associated in pathological processes such breast cancer, endometrial cancer, and osteoporosis (NCBI, 2021). ER acts as a ligand-dependent transcription factor, and ligand binding to the LBD results to activation of gene transcription, leading in the progression of breast cancer (Takeshima et al., 2020). Of all diagnosed breast cancers, about 70%-80% are estrogen receptor (ER)-positive breast cancers (Piscuoglio et al., 2018), hence the most common subtype of breast cancer. Targeting and depriving the ER pathway is a major and foremost therapeutic strategy to manage metastatic and early-stage disease (Reinert et al., 2017).

TP53 is heavily associated with cancers as it encodes a protein called p53 that is known to suppress tumor by facilitating response to diverse cellular stresses such as apoptosis, senescence, cell cycle arrest, DNA repair or changes in metabolism, as well as negative regulation of cell division to regulate expression of target genes (Gene Cards, n.d.). However, this gene is also known as one of the most observed mutated genes in breast cancer observed in 30-35% of all cases acting as a factor for a negative prognosis. Moreover, high prevalence of mutated tp53 in tumors such that in breast cancer are likely to be aggressive as frequently reported in approximately 80% of triple negative breast cancer cases, resistant to chemotherapy and radiotherapy prognosis (Duffy et al., 2018). With this, tp53 shows great potential as a target and biomarker in the development of various therapeutics for breast cancer patients but should be of greatest consideration as the role of tp53 in breast cancer management remains unclear.

Berberine as one of the topmost bioactive compound in Coix has been reported by various studies to possess anti-tumor activity. It has been a hot topic in experimental research in recent years and of great consideration for use in breast cancer adjuvant therapy as it can exhibit potent anticancer activities through its ability to inhibit cell cycle and induce apoptosis of cancer cells (Magalhães et al., 2021). Furthermore, Berberine can impede tumor growth and metastasis of triple negative breast cancer cells by suppressing the expression of transforming growth factor beta 1 (TGF-B1), a factor involved with poor prognosis on breast cancer (Magalhães et al., 2021). Studies then reveal that Berberine appears to be good candidate as chemosensitizer and chemotherapeutic in treating breast cancer offering improved therapeutic efficacy and safety

Beta-sitosterol is one of the several and principal phytosterols in plants with a chemical structure similar to that of cholesterol (Rashed 2020; Van, 2000). Beta-sitosterol has been reported to have antidiabetic activity by increasing the fasting plasma insulin levels, decreasing fasting glycemia, and reducing glucose levels (Saeidnia, 2014) and anticancer activity including breast cancer, prostate cancer, colon cancer, lung cancer, stomach cancer, and ovarian cancer (Rashed, 2020). Studies suggested that Beta-sitosterol is an effective apoptosis-promoting agent and that integration in the diet could provide a preventive measure for breast cancer (Awad et al., 2007).

A significant reduction in the degree that fresh drug candidates are being transformed into clinical treatments with a high efficacy rate and value has been evident over the last few decades (Hopkins, 2008). With the breakthrough and rapid advancement of fields of bioinformatics, systems biology, and polypharmacology (Li & Zhang, 2013; Shi et al., 2014; Zhang et al., 2019), the concept of network pharmacology was presented by Andrew L. Hopkins (Shi et al., 2014) in order to examine the effect and intermediation of drugs as well as to demonstrate the synergistic principle of multicomponent drugs to search for multiple target treatment having high efficacy and low toxicity activity rather than single target (Tang et al., 2016; Zhang et al., 2013). Moreover, network pharmacology, by means of a systematic approach, examines on the interactions between drugs, targets, diseases, and compound-proteins/genes-disease and visually displays the network of drug targets. Still, it explored the influence of drugs on the biological network from a holistic perspective and is able to explain complexities among biological systems, drugs, and diseases from a network perspective (Zeng & Yang, 2017; Lui & Bai, 2020; Zhang et al., 2019). Consequently, network pharmacology is of great help in identifying the mechanism of action and assess the efficacy of drug through discovering the influence of compounds to the biological network (Gu et al., 2013).

Network pharmacology highlights the change of paradigm from the traditional “one target, one drug” to “network target, multicomponent therapeutics,” drug discovery (Li et al., 2014). According to Zhang et al. (2013), the advantages of network pharmacology study involve: (1) regulation of the signaling pathway with multiple channels; (2) increase in the efficacy of drug; (3) decrease of side effects; (4) increase in the success rate of clinical trials and; (5) reduction in the financial costs in drug discovery. As a result, this network-based drug discovery is deemed a promising method and has great potential to become the next generation approach of drug discovery (Li et al., 2014).

With the rapid advancement of research, network pharmacology, including the investigation of traditional medicines has become a new and prevalent method for drug mechanism research and drug development (Zhang et al., 2019; Lee et al., 2018). Numerous efforts have been made to employ the method to examine the complex of ingredients, unknown targets, as well as pharmacological mechanisms of herbal formulas to elucidate and provide understanding into the complex mechanisms of herbal formulas used to treat complicated diseases (Lee et al., 2018).

Through scientific exploration, the elucidation of various biological processes of *Coix lacryma-jobi* has been established despite the existence of its therapeutic application for thousands of years. Hence, it paved way for the occurrence of integration of traditional and modern medicine (Kuo, 2012) which was also evident from the different bioinformatics tools. Traditionally*, Coix lacryma-jobi* seeds have been utilized as herbal medicine for the treatment of diabetes, asthma, inflammation, cough, diarrhea (Choi et al., 2015), pulmonary edema, wet pleurisy, chronic ulcer, dysuria (Feng et al., 2020), rheumatism, osteoporosis, (Zhang et al., 2019) chronic gastrointestinal disease (Choi et al., 2015; Feng et al., 2020), and as a diuretic (Zhang et al., 2019, Zhu, 2017; Wu et al., 2014).

*Coix lacryma-jobi* has long been utilized in traditional Chinese medicine for several diseases particularly cancer (Woo et al., 2007) and has been reported to possess anti-cancer properties (Xi et al., 2016). Studies have also proposed that sterols, one of the constituents of *Coix lacryma-jobi,* have a protective role from various types of cancers, including breast cancer (Chang et al., 2018). A pharmaceutical micro-emulsified injectable drug obtained from coix seed oil, called Kanglaite Injection (KLT), showed anticancer activity and was approved by the Chinese Ministry of Public Health to treat various tumors and common types of cancers such as breast, lung, liver, and pancreatic, and gastric cancer. The treatment was clinically approved in China, Russia, and USA Food and Drug Administration (Fang et al., 2020; Woo et al., 2007; Yu et al., 2008).

Despite evidences supported by clinical data on the usage of coix extract preparation in the treatment of cancer, particularly breast cancer, there has been insufficient studies that investigate and establish the biological basis and mechanisms of its activity (Woo et al., 2007). Our study had elucidated on the pharmacological underpinnings on the potency of adlay against breast cancer. It is hoped that this study may serve as a reference and guidance in the drug development for breast cancer treatment.

## CONCLUSION

Our study provides the pharmacological basis elucidating on the potency of *Coix lacryma-jobi* against breast cancer using a network-based approach focusing on its bioactive components, gene targets and pathway analysis. There were 201 compounds in *Coix lacryma-jobi* identified as bioactive compounds that target numerous breast cancer-associated genes. Based on the Enrichment Analysis, *Coix lacryma-jobi* targets were enriched in GO terms mostly associated with positive regulation of transcription from RNA polymerase II promoter, drug response, steroid hormone receptor activity, and protein binding. With the efficacy of *Coix lacryma-jobi* for breast cancer treatment, it is noteworthy that it may have similar impacts in treating other types of cancers. Nonetheless, its subsequent drug development will be worth a step forward for a breast cancer-free society.

**Table 1.**
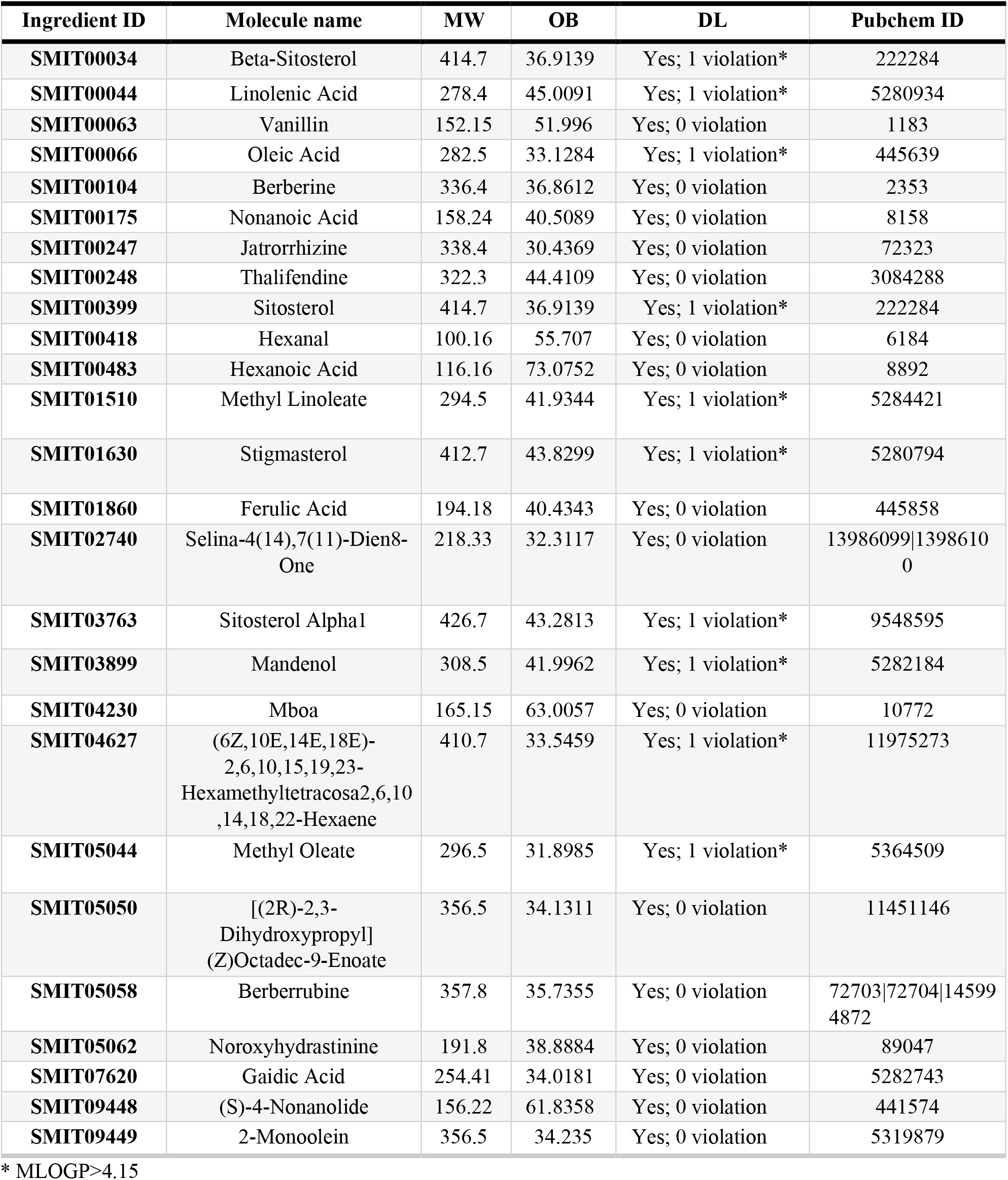
A list of selected compounds of *Coix lacryma-jobi* for network analysis.

## Acknowledgements

The authors are grateful to the unwavering support of the Biology Department faculty of the Polytechnic University of the Philippines. Special mention goes to Prof. Lourdes Alvarez, Prof. Julie Charmain Bonifacio, and Prof. Chester C. Deocaris for their guidance, patience, and moral support.

## Author Contribution

MVA and CCD conceptualized & designed the study, interpreted the data, and edited the manuscript; AT, AMF, and JRD performed data mining, analysis, and wrote the draft manuscript. AB performed data analysis, wrote the draft, and edited the manuscript. All authors read and approved the final manuscript.

## Data Availability

The data supporting the findings of this study are available in the paper and from the corresponding author, Malona V. Alinsug, upon request.

## Ethics Approval

NA

## Consent to Publish

All authors read and approved the final manuscript.

---

**Supplementary Table S1.**
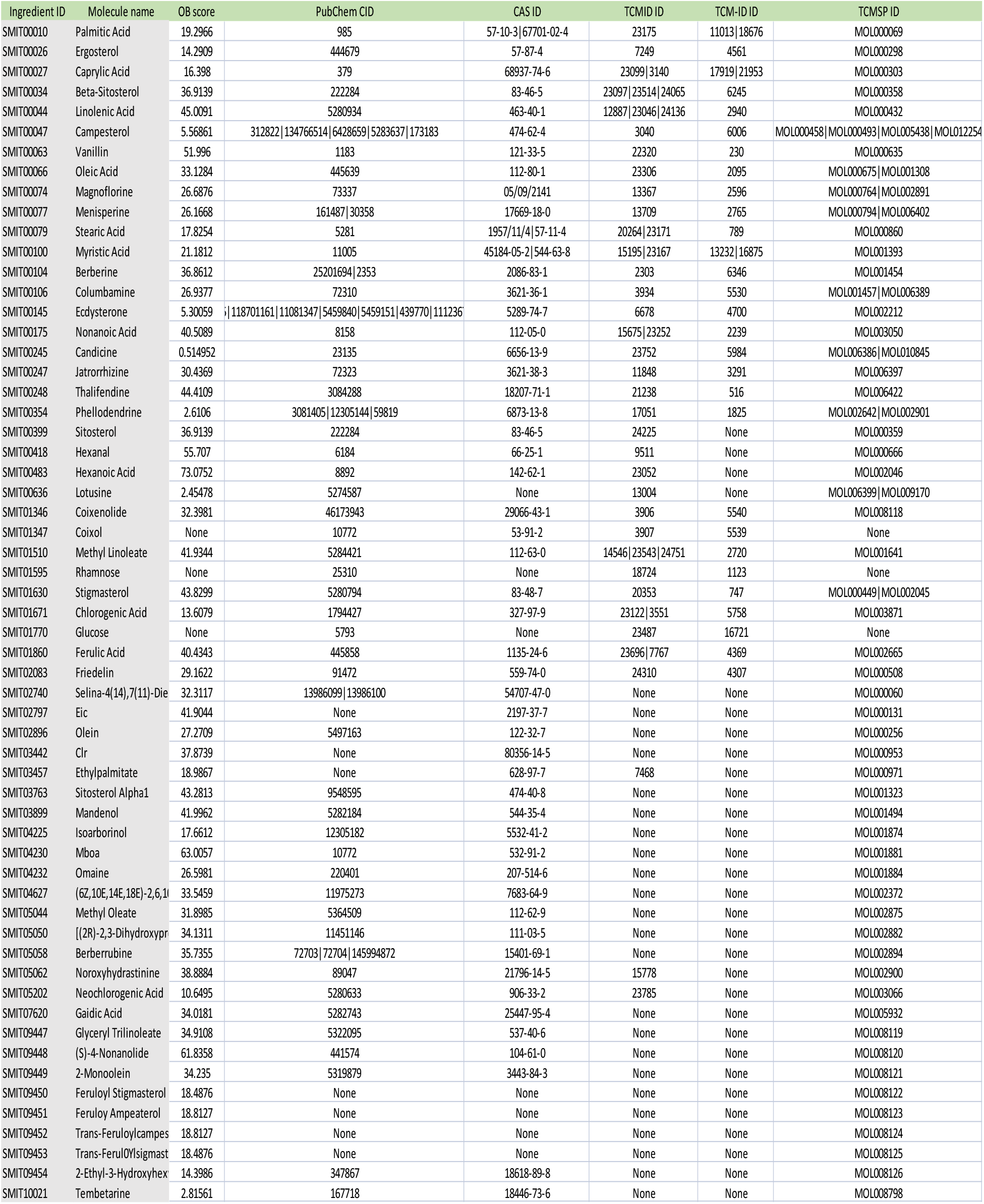
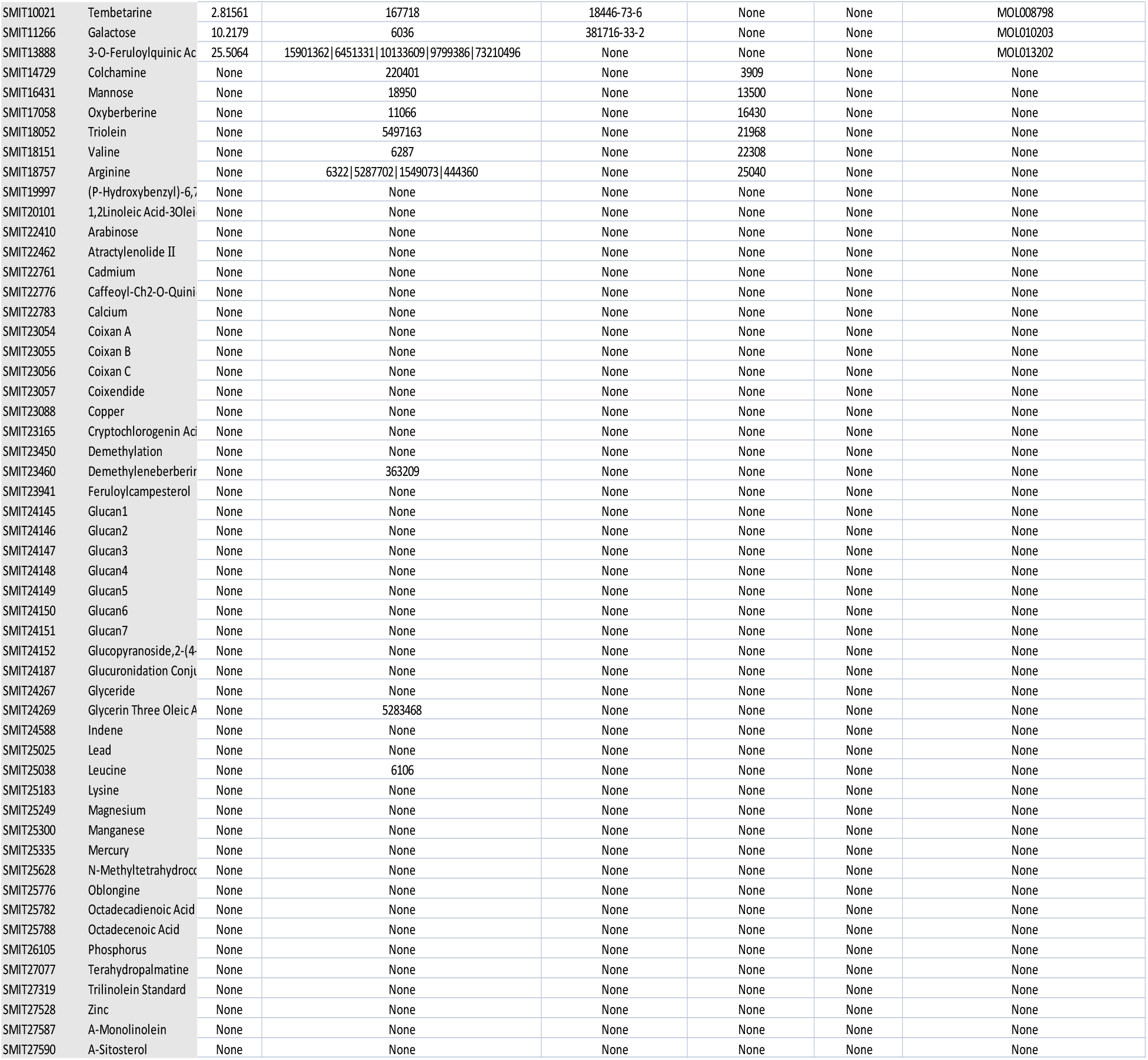

**Supplementary Table S2.**
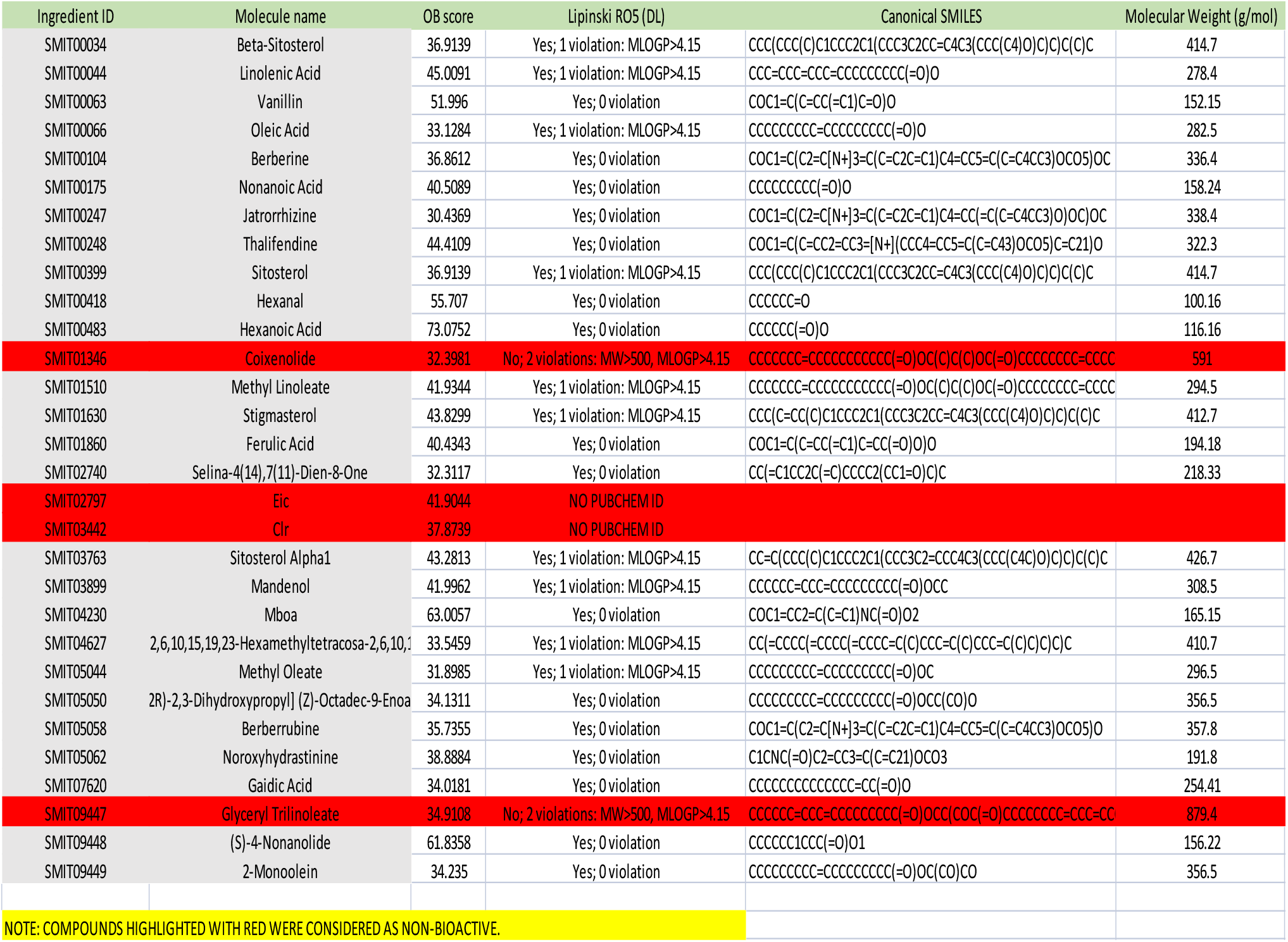

